# Human-Robot Interaction with Robust Prediction of Movement Intention Surpasses Manual Control

**DOI:** 10.1101/2020.12.09.416735

**Authors:** Sebastijan Veselic, Claudio Zito, Dario Farina

## Abstract

Designing robotic assistance devices for manipulation tasks is challenging. This work aims at improving accuracy and usability of physical human-robot interaction (pHRI) where a user interacts with a physical robotic device (e.g., a human operated manipulator or exoskeleton) by transmitting signals which need to be interpreted by the machine. Typically these signals are used as an open-loop control, but this approach has several limitations such as low take-up and high cognitive burden for the user. In contrast, a control framework is proposed that can respond robustly and efficiently to intentions of a user by reacting proactively to their commands. The key insight is to include context- and user-awareness in the controller, improving decision making on how to assist the user. Context-awareness is achieved by creating a set of candidate grasp targets and reach-to grasp trajectories in a cluttered scene. User-awareness is implemented as a linear time-variant feedback controller (TV-LQR) over the generated trajectories to facilitate the motion towards the most likely intention of a user. The system also dynamically recovers from incorrect predictions. Experimental results in a virtual environment of two degrees of freedom control show the capability of this approach to outperform manual control. By robustly predicting the user’s intention, the proposed controller allows the subject to achieve superhuman performance in terms of accuracy and thereby usability.

## 1 Introduction

Automation is leading to major societal changes, with an estimated 50% of current jobs to be subjected to automation. However, automation often requires intelligent semi-autonomous robots operated by human users with physical human-robot interaction (pHRI) (*4*). For example, there is already a growing need for efficient controllers for robot manipulators involved in operations such as nuclear waste disposal or manufactoring processes (*17*).

Furthermore, the progressive ageing of the world population (*7*) will increase the need for robotic assistants that will be teleoperated by human users in cases such as, for example, surgical procedures (*23*). For these devices, algorithms to predict the user intention are a crucial component of the control system. In both examples of pHRI in automation and assistive devices, the purpose of pHRI is to achieve the user’s planned goals in a given domain (*16*). Here, planned goals are defined as motor command sequences to be executed to achieve a final state (*x_T_*) from an initial one (*x*_0_).

However, controller systems employed in pHRI settings will need to go above and beyond being able to only infer movement intention and helping achieve planned goals. Additionally, they will need to i) send feedback to users (*16*) and ii) adjust their own functioning accordingly after inferring intention (*28*). Furthermore, there will be a need to actively infer long-term goals from sequential data points (*19*), and learn goals from continuous corrective feedback designed to teach them about planned goals (*1*). All these characteristics of pHRI represent areas of ongoing research (*1, 8, 19, 28*). On a conceptual level, Fig. 1 summarises the relevant components of pHRI for our work. The user sends an intention which is intercepted by a controller. The controller decodes that intention in a given task and sends (corrective) feedback to the user. Here, the intention may be decoded from myoelectric signals generated by the muscles of the user while performing the task or from the kinematics of the user that encodes the intention of moving to a particular location. The controller and the user jointly influence the interaction with the environment when their actions are modulated according to an arbitration process (*16*).

**Figure 1:**
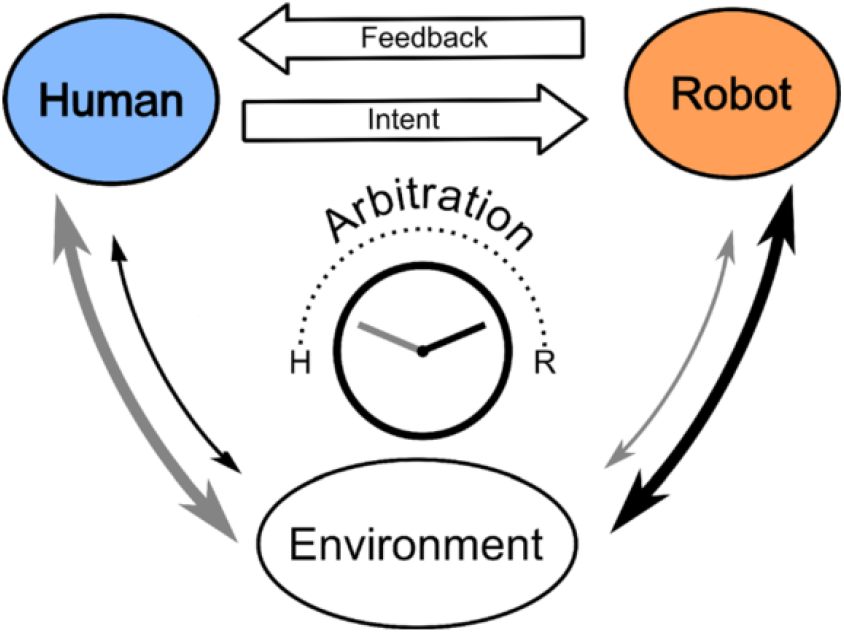
Adapted from (*16*). A scheme of the interplay between robot feedback and human intention in pHRI. Humans receive feedback from the robot (controller) system once it decodes human task-specific intention. The robot and the human jointly exert control on the environment where the degree of control from either is determined through arbitration for a given domain.

In this work, a novel way of achieving this pHRI coupling was developed by enhancing the robot component with context- and user-awareness. Context-awareness enables the system to perceive the environment and to identify potential actions for the user. User-awareness enables the system to recognise the user intent and to assist the motion. The central contribution of this work is achieving high accuracy in reaching tasks by including user- and context awareness in a pHRI system.

Context-awareness was enabled by manual construction of a set of candidate grasp targets and their reach-to-grasp trajectories for the testing scenario representing a scene with multiple graspable objects. The set of candidate trajectories represents the optimal trajectories to reach these target states (i.e. graspable objects) from a starting state and are the input to the controller. In this work, the robot manipulator is dened as a robot arm and a gripper. Thus, each trajectory is defined as a set of robot’s configurations for both the arm and the gripper. This manual construction of optimal trajectories can be substituted by automatic determination of the trajectories, as demonstrated in recent advances in autonomous robotics for grasping and manipulation of novel objects (*11, 32, 33*). For example, Kopicki and Zito proposed generative models to learn and transfer manipulative skills in robotics (*11, 21, 32–34*) where a set of contact models for a particular grasp are learned from a single demonstration. Each of these models learns a set of local features describing the contacts between the object and the robot’s end effector, so that it can generate a set of potential grasp candidates for a novel object by seeking for similar local features on the novel object’s surface. In (*32, 33*), these models have been extended for planning reach-to-grasp trajectories to deliver the robot’s end effector to the target grasp even in case of uncertainty due to an incomplete representation of the objects, i.e. a partial point cloud representation.

User awareness was implemented through a time-variant LQR controller (*12, 14*) (TVLQR) that filters the motion commands of a user at each time step and assists the user along the trajectories of the candidate grasps, thereby inferring movement intention. In control theory, a conventional LQR algorithm uses linearisation of the dynamics of the motion around a nominal trajectory to compute an optimal feedback control law for the system (*12*). For complex nonlinear systems, it is not possible to linearly approximate the entire dynamics of the system, thus the time-variant variant of the algorithm produces a set of locally optimal feedback control laws around a set of pre-determined waypoints along the trajectory. The controller proposed in this study extends the TV-LQR to deal with several trajectories at the same time. For each waypoint belonging to any of the nominal trajectories, we compute the feedback controller to either follow the trajectory or moving towards a neighbouring one. The proper feedback control law is selected online by filtering the user’s motion input. Thus, in this work, user assistance is defined as support enabling the user to stay close to the optimal trajectory for the expected target state or grasp by sending corrective feedback. If the user recognises that the system is moving towards an incorrect target, the user is expected to apply a corrective motion that we use to compute the feedback control to guide the system towards a new best candidate. The TV-LQR is ideally suited for such computation as it can deal with potential changes in system dynamics and user input by computing optimal corrective feedback for all points on a given trajectory (*14*), without the need of pre-defined thresholds for the switching conditions. Furthermore, the gripper’s orientation and the fingers configurations are inferred from the pre-computed trajectories, allowing the user to focus only on how to guide the arm towards the desired candidates. Grasping motions to align the gripper with the target object and the fingers congurations are therefore not directly controlled by the user, but executed along the users chosen trajectory.

The proposed approach differs compared to other existing methods that mostly rely on crude force feedback via a haptic device to drive the user towards a pre-defined target pose or desired trajectory, e.g (*2, 20*). The approach proposed here employs a predict-then-blend approach, in which the most likely intention of the user is estimated first and then assistance is provided in the relative manipulation task that was tested in simulation. Previous work related to intention detection and user assistance has been usually approached with myoelectric or EEG recordings which have limitations associated with low signal to noise ratio (reviewed in (*16*) (*15*)). Conversely, the proposal here is that motor intention can be estimated directly from the user kinematics, representing their final desired output. Namely, it has been previously shown that the LQR controller is effective at solving the problem of user assistance (*3*) and correcting user-given input (*18*) (*13*). Furthermore, existing work aiming to predict user intention is based on computing the distance between the user’s current configuration and the desired one (*20*). However, such an approach cannot scale to complex and dynamic environments or tasks because it ignores the history information and kinematic limitations of the teleoperated robot. Moreover, while it has been previously proposed that predictions could be based on learning from demonstration (*5, 9, 31*), these methods need to learn the user’s intention from complex motions and plan goal-dependent trajectories online to adapt to novel contexts. Contrary to this, the proposal presented here does not require learning a complex task-dependent function for the operator. Furthermore, the controller is constructed online and easily adapts to erroneous initial predictions enabling smooth transitions towards the next most likely target.

In addition to predicting user intention, shared control involves using the predictions to support the user in achieving the expected goal. Existing methods propose switching between manual and autonomous mode depending on the confidence level of the predicted goal. That is, when confidence of achieving the goal is low, the system behaves as an autonomous controller (*6*). However, thresholds employed in such methods require adjustments due to the tasks at hand, and a constant tracking of the confidence level to enable switching back to manual control in case of unexpected occurrences. More sophisticated methods use virtual fixtures (*22*) to compare the user’s input to the desired motion and provide some sort of feedback. Here, a more advanced approach that enables soft integration of the input commands and the predictions is proposed. The TV-LQR estimates the most likely input. If the user matches the expectation, the commands are filtered to follow the expected trajectory. Otherwise, if the user is moving away from the expected direction, the feedback controller automatically dilutes its adjustments and enables the user to smoothly transition to the closest expected motions which sits in the region of space where the user is currently moving towards.

In summary, for the first time, a generic formulation for user’s motion intention detection based on the use of the time-varying version of LQR to filter and optimise human motor control in a grasping scenario is proposed. Moreover, using a predictive extension of the LQR the desired goal of the user is predicted and inferred in an unsupervised manner. Finally, the controller is tested and compared to manual control, as an important aspect of pHRI will be superhuman performance according to at least one objective function (e.g., accuracy, computational cost, or speed) (*30*) which will facilitate the adoption of these approaches in practice.

## 2 Results

The controller was first tested in simulation through characterisation runs to determine the optimal parameter set (*τ*, *α*, *S*) for the testing domain and to investigate their impact on accuracy (see Sec. 4). Here, accuracy was defined as the Euclidean distance between the final system pose and the grasp target pose for a specific target trajectory after 50, 100, and 200 trajectory waypoints under constant input. The testing scenario was designed to mimic real-world usecases where such controllers will be used (e.g. nuclear waste disposal and similar cases where a human teleoperates a robot arm). To make the results more robust and generalizable to the real world, the parameter combinations were investigated at different trajectory lengths that could be dynamically observed in real world settings.

The goal of each execution was to provide constant input (i.e. velocity) in *x*, *y*, and *z* coordinates to drive the controller from a starting pose towards a final pose. The final pose was always an arbitrarily picked grasp target pose of a specific trajectory that remained the same throughout an execution run. By keeping the input and final pose constant, the contribution of individual parameters (*τ*, *α*, *S*) on accuracy was determined. Crucially, despite providing constant input and a constant final pose, non-additive unit noise (Eq. 14) was added at each waypoint for each execution run and thus all reported results include noise because the aim was to design a robust control system that would generalize to real world domains.

The optimal parameter set was then used on human subjects where the controller was directly compared to manual control with a within subjects design. Due to a within-subjects design, the sample size was (*N* = 6) subjects and is comparable to previous related work (*25, 26, 29*).

### 2.1 Characterisation runs

#### 2.1.1 Trajectories with 50 waypoints

The characterisation run results with 50 waypoints are reported first (Supplementary Materials Fig. 3a). Two independent-sample Welch t-tests showed that the parameter setting *α* = 0.5 (*M* = 9.20 ± 0.33) yielded best accuracy compared to both *α* = 0.1 (*M* = 11.62 ± 0.23, *t*(3427.5) = 244.45, 95% CI [2.40, 2.44], *p* = 2.2*e*-16), and *α* = 0.9 (*M* = 11.59 ± 0.26, *t*(3206.2) = 258.72, 95% CI [2.38, 2.42], *p* = 2.2*e*-16). In Fig. 3a it was also revealed that different parameter values of *S* and *τ* did not affect accuracy, irrespective of which combination was used during the characterization runs. This is also reflected by independent-sample Welch ttests where the pooled *S* = 1 with the *S* = 5 (*t*(3598) = 0.21, 95% CI [−0.07, 0.09], *p* = 0.83), and *S* = 9 (*t*(3597) = 0.05, 95% CI [−0.08, 0.08], *p* = 0.96) across all *α* and *τ* values were compared and did not show a difference in accuracy. Similarly, when the same analysis was repeated to investigate the effect of *τ* on accuracy, the results showed that the pooled *τ* = 0.1 was comparable to *τ* = 0.5 (*t*(3597.9) = 0.22, 95% CI [−0.07, 0.09], *p* = 0.83), and to *τ* = 0.9 (*t*(3597.9) = 0.11, 95% CI [−0.07, 0.08], *p* = 0.91) across all *α* and *S* values. In sum,in trajectories with 50 waypoints, only the *α* parameter had an impact on accuracy.

**Figure 2:**
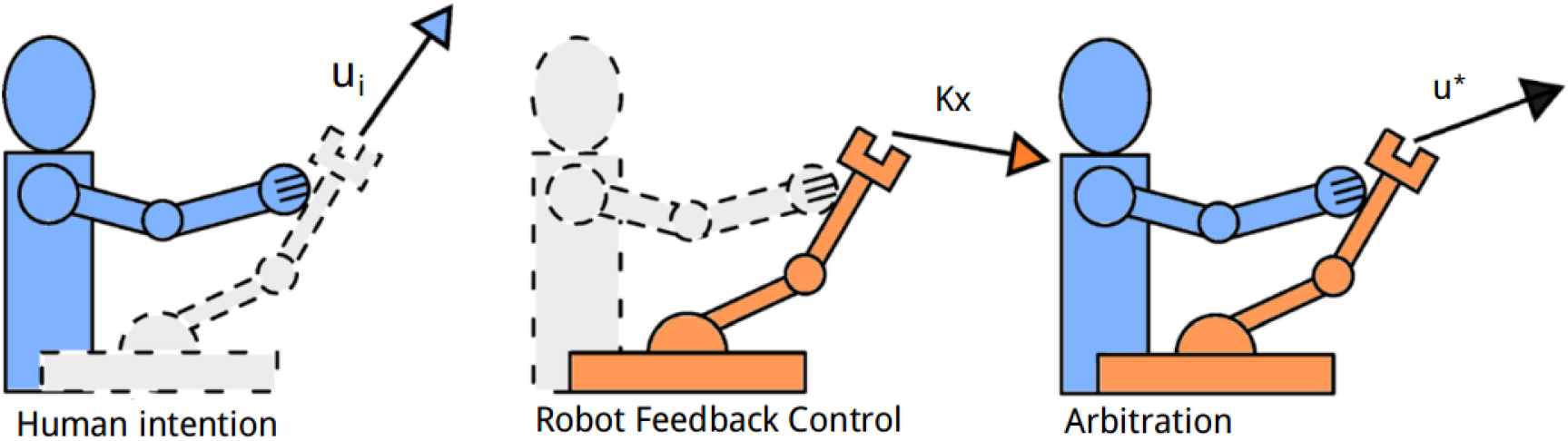
Adapted from (*16*). A sample example of the model obtained above catered for our approach (see Sec. 4). The final exerted control signal (*u*^∗^) exerted is a combination between the user input (*u_i_*) and the controller feedback control, which is in our case determined by the *K* matrix. On the figure it is represented by *Kx* to denote its impact on the system state (*x*) as well, to include all relevant terms used when computing *u*^∗^.

**Figure 3:**
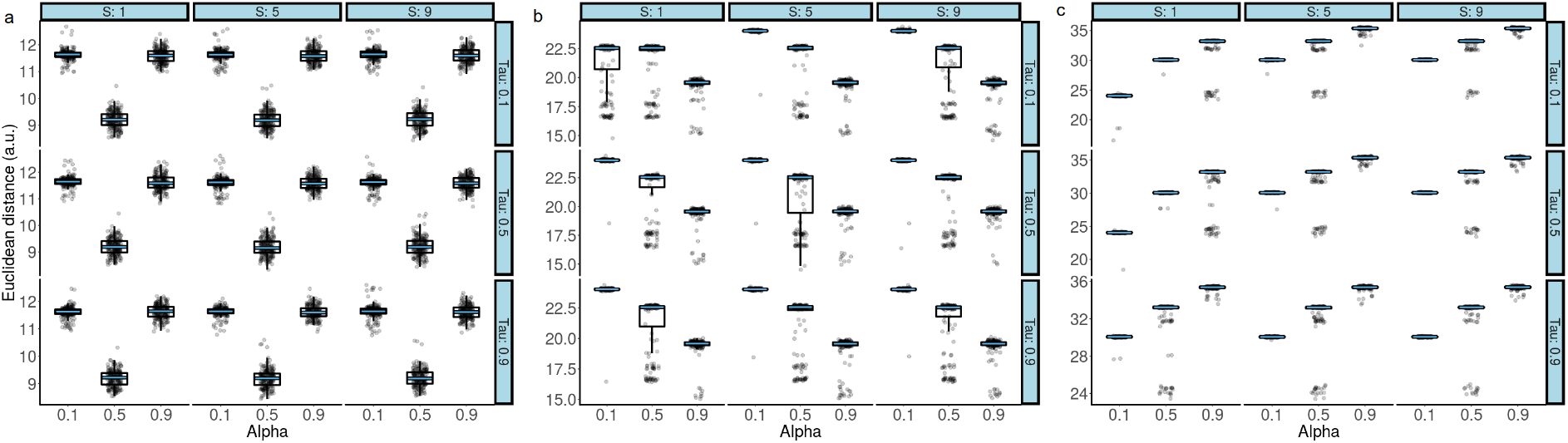
Boxplots overlaid with raw data showing the effect of (*τ*, *α*, *S*) on the Euclidean distance after passing 50 trajectory waypoints (a), 100 trajectory waypoints (b), and 200 trajectory waypoints (c). The black line of the box plot denotes the median value. The upper and lower bound of the boxplot denotes the 1st and 3rd quartile of the dataset. The boxplot whiskers denote the 1.5*IQR. The superimposed blue line on the boxplots denotes the median and has been added for clarity purposes. In all three panels, the *α* parameter had the strongest effect on Euclidean distance

#### 2.1.2 Trajectories with 100 waypoints

The same set of analyses was applied to trajectories with lengths of 100 waypoints. In contrast to the results from Fig. 3a, Fig. 3b reveals that *α* = 0.9 (*M* = 19.31 ± 0.99) led to the lowest Euclidean distance in all conditions, i.e. had the best accuracy. To statistically evaluate this, Wilcoxon rank sum tests were used and the difference between different levels of the *α* parameter was assessed. Non-parametric tests were used because of bimodal distributions that were present in the majority of conditions on Fig 3b. Again, both *S* and *τ* were pooled to obtain means corresponding to *M* = 23.69 ± 1.23, and *M* = 21.35 ± 2.26, respectively. A comparison of *α* = 0.9 to *α* = 0.5 (*W* = 673730, 95% CI [2.93, 2.96], *p* = 2.2*e*-16), and *α* = 0.1 (*W* = 92510, 95% CI [1.45, 1.47], *p* = 2.2*e*-16) showed that the difference observed in Fig. 3b was statistically significant.

#### 2.1.3 Trajectories with 200 waypoints

In the last set of characterisation runs, the lowest Euclidean distance between the final system pose and the target grasp pose (i.e. the highest accuracy) was obtained when *α* = 0.1. Thus, when the strongest corrective feedback was given by the TV-LQR controller (i.e. *α* = 0.1), the smallest discrepancy between the final system pose and the target grasp pose was observed. As in the case of previous trajectory lengths, when means were pooled across *S* and *τ*, it was observed that *M* = 28.71 ± 2.55, *M* = 31.81 ± 2.41, and *M* = 34.59 ± 1.94, for *α* = 0.1, *α* = 0.5 (*W* = 466280, 95% CI [3.15, 3.16], *p* = 2.2*e*-16), and *α* = 0.9 (*W* = 75702, 95% CI [5.315.33], *p* = 2.2*e*-16), respectively. That is, both statistical tests showed that *α* = 0.1 had the the best accuracy. Notably, Fig. 3c also revealed comparable performance in the case where *τ* = 0.5 and *τ* = 0.1 (*W* = 1589900, 95% CI [−0.02, 0.01], *p* = 0.33), but improved performance when compared to *τ* = 0.9 (*W* = 1307800, 95% CI [0.07, 0.16], *p* = 2.2*e*-16).

Overall, the best accuracy was observed in trajectories with the smallest number of way-points (50) and equal weighting (*α* = 0.5; Fig. 3a, column with *α* = 0.5). Increased trajectory lengths led to additional variation (Fig. 3b and 3c).

When the trajectory length was increased to 200 waypoints, the lowest Euclidean distance was observed with *α* = 0.1. Equations 7, 11, and 12 (see Sec. 4), reveal this to be a sensible result because the total error can be minimised over time by maximising the adherence to optimal corrective feedback given by the TV-LQR. This is exactly what the given parameter setting achieved within the parameter space of *α* values measured here.

### 2.2 Comparing manual and assisted control

With the obtained optimal parameter set, the controller was tested on (*N* = 6) subjects. In this section, the terms manual control and manual condition are used interchangeably. Similarly, the terms shared or assisted control/condition are used interchangeably as well. The main hypothesis was that shared control would improve accuracy compared to manual control. Crucially, we hypothesized that this would be observed for each subject, as the improvement in accuracy would need to be relative to a subject’s own performance, as opposed to only an improvement in accuracy across subjects.

Before starting the task, subjects were told they would be providing input to guide a controller in a simulated scene with several possible target grasps by moving a computer mouse. By increasing or decreasing the position of the mouse along the *x* and *y* axis in physical space, they would be increasing or decreasing velocity for the respective dimension in the simulation, while the *z* axis would be slowly increasing to mimic moving from a starting system pose towards target grasp poses. They were also told that there was always an optimal trajectory from their starting position to any of the target grasps they could see, which the TV-LQR would try to keep them close to in one of the conditions. Finally, they were presented with information of their current position in terms of *x, y*, and *z* coordinates, their velocity, the coordinates of several final target grasp poses, the waypoints of the optimal trajectories closest to their current position, and the coordinates of the trajectory they were actually closest to, in case it was not one of the target ones.

The subjects’ task was to pick one of three, arbitrarily defined, target grasp poses and provide input to move the controller such that they would achieve a final system pose that would be as close as possible to their target grasp pose (i.e. the final pose of a specific trajectory) (Supplementary Materials Fig. 4). They were required to perform this task either without assistance or with assistance by the TV-LQR controller.

Furthermore, in half of the trials, subjects were required to switch their targeted grasp halfway through the execution (switch condition) as opposed to the other half of trials where this was not necessary (non-switch). This factor was added to evaluate the robustness of the proposed controller design. The goal of the switch condition was to mimic users who would determine throughout the execution of a movement that their desired goal is elsewhere and would therefore need to switch what they previously intended on doing. Importantly, the main manipulation of the design was the inclusion of the assisted and manual condition which were orthogonal to the switch and non-switch condition. All subjects were tested on trajectories with a length of 200 waypoints. Three sources of noise were added to this part of the experiment to further stress the controller. These included noisy trajectories and noisy system updates at the level of each waypoint and for each subject (Supplementary Materials Fig. 5). Finally, to ensure the controller design was intuitive enough such that naive subjects would be able to comprehend it without substantial training, most subjects were previously unfamiliar with the controller, the design, and the task. Through this, a possible confound of training and over-trained subjects that could have affected these results was eliminated.

#### 2.2.1 Shared control improves accuracy across subjects in the switch and no switch condition separately

To investigate whether shared control improved accuracy, the distance in the shared and manual condition across subjects was compared. This comparison revealed that the shared control (*M* = 24.26 ± 3.46) condition, where TV-LQR was providing assistance, led to statistically significantly less error compared to the manual condition (*M* = 27.48 ± 4.85, *W* = 106430, 95% CI [2.31, 3.18], *p* = 2.2*e*-16, Fig. 4c). To further investigate whether this was true in both the switch and non-switch condition, data was split according to both and compared (Fig. 4ab). In both the switch and non-switch condition the results remained the same, i.e. the assisted condition led to higher accuracy, or lower error, compared to manual control: switch condition (shared -*M* = 24.33 ± 3.51; manual -*M* = 27.26 ± 4.56, *W* = 27858, 95% CI [1.98, 3.17], *p* = 6.811*e*-16), non-switch condition (shared -*M* = 24.20 ± 3.41, manual: *M* = 27.69 ± 5.11, *W* = 25444, 95% CI [2.30, 3.55], *p* = 2.2*e*-16). These results confirmed that the employed TV-LQR controller outperformed manual control.

**Figure 4:**
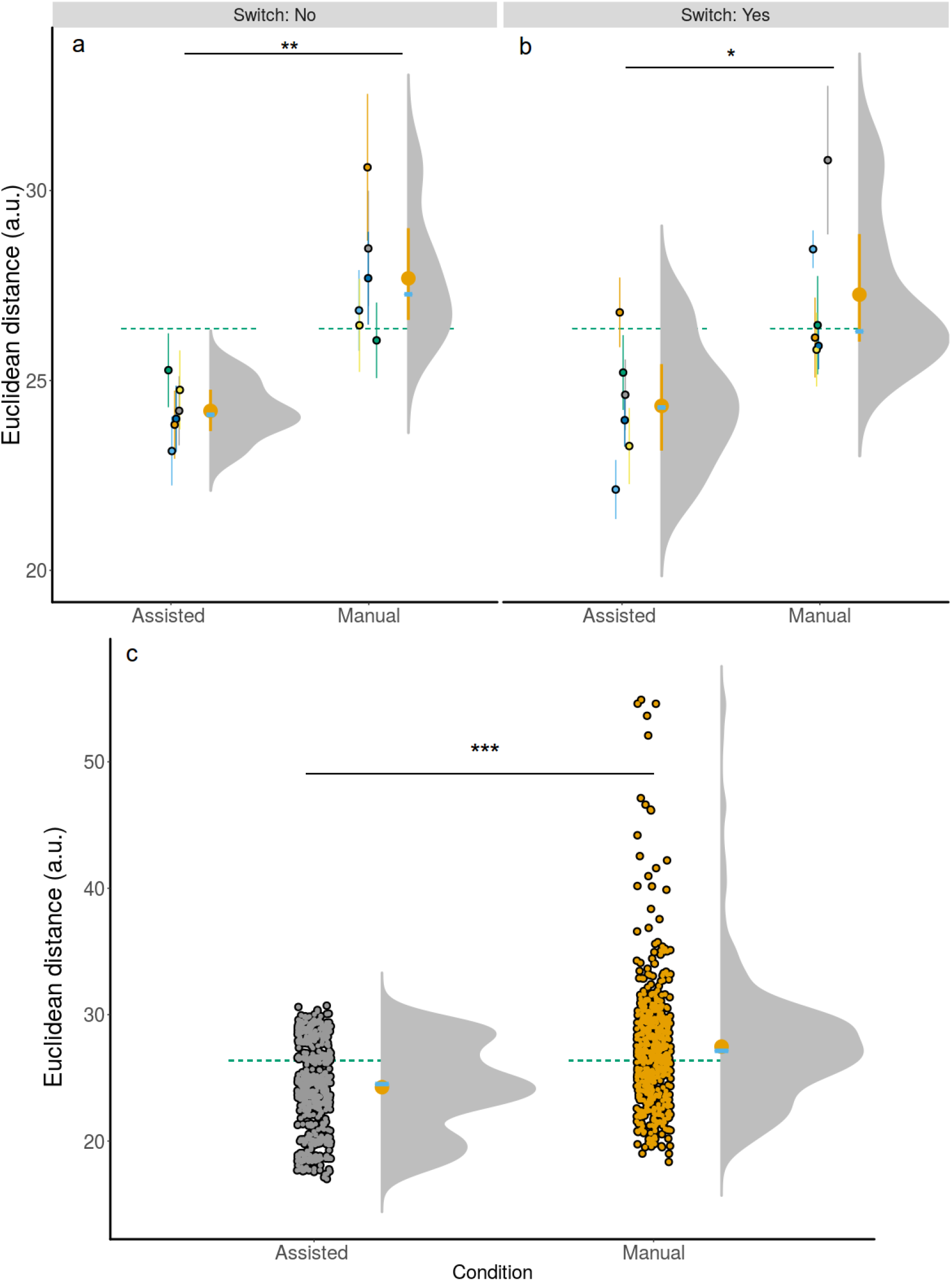
Accuracy of individual conditions for all subjects. **a)** and **b)** show subject color-coded summarised information (*N* = 6). The accompanying lines denote the 95% CI for the non-switch **a)** and switch **b)** condition. **c)** shows the same information collapsed across both **a)** and **b)** in addition to showing raw trial information across all subjects to show the full distribution across all trials. The green dotted line in **a)**, **b)**, and **c)** is the grand mean collapsed across all conditions. The yellow error bar corresponds to the condition mean and 95% CI. The superimposed blue bar shows the condition median. *p* < .001: ***, *p* < .01: **, *p* < .05: *. All three panels show that the assisted condition resulted in higher accuracy compared to manual control.

Because the previous comparisons were performed on pooled data and because the observations were not independent, a more rigorous set of analyses was performed where the the performance across trials was summarised by subject and by condition (assisted and manual). This enabled performing further paired, two-sided Welch t-tests that revealed comparable results. That is, subjects improved their performance in the assisted condition (*t*(11) = 4.69, 95% CI [1.70, 4.72], *p<.*001) compared to the manual one. Moreover, this difference was more apparent in the non-switch (*t*(5) = 4.07, 95% CI [1.29, 5.69], *p<.*005) compared to the switch (*t*(5) = 2.57, 95% CI [0.01, 5.86], *p<.*05) condition.

Finally, when between-subject variance was accounted for and modelled by linear-mixed models (LMM), a similar result to the one reported above was observed. An LMM with the condition (assisted, manual) and information about switching (Yes, No) as fixed factors, subjects as random intercepts with varying slopes for both condition and switching information, and trials as random intercepts to predict the observed Euclidean distance was fit. A Type III Analysis of Variance with Satterthwaite’s method showed that condition (*F* (5.00, 1) = 22.91, *p<.*005) but not switching information (*F* (5.67, 1) = 0.14, *p* = 0.72) nor their interaction (*F* (1093.16, 1) = 1.45, *p* = 0.23) were significant predictors for the final observed distance. Individual t-tests approximated using Satterthwaite’s method for condition showed that a change from shared to manual resulted in a greater Euclidean distance between the optimal and actual target state (*β* = 3.49 ±0.71, *t*(6.28) = 4.92, *p<.*005). This implies an increase in the Euclidean distance between the final system pose and the final grasp target pose by *β* = 3.49 due to performing manual, as opposed to shared control. In other words, it implies a deterioration in accuracy in the manual compared to the shared condition, in line with our previous results.

#### 2.2.2 Assisted control improves accuracy in each subject

As the most stringent criterion for determining whether the controller was better compared to manual control, the conditions were compared for each individual separately. Namely, the focus of this work was to see whether for each subject individually, enabling shared control on a task would yield an improvement in their accuracy compared to manual control where they received no assistance. To test this, we assessed whether the regression lines for each subject showed a positive slope (*β*) from the assisted to the manual condition. A positive slope would in this case mean that there was a deterioration in performance when going from assisted to the manual condition and thereby a general improvement under the TV-LQR assisted condition. This means that if all subjects individually exhibited a positive slope, then the observed effect would not driven by single outliers but all observed datapoints (i.e. all subjects). Indeed, this is what was observed (Fig. 5-6). The *β*s for individual subjects: *M* = 3.21, *Min* = 1.02, *Max* = 5.23 were positive and none had a negative beta. These results indicate that accuracy improved when they received TV-LQR assistance in the shared control condition and are in line with the hypothesis. The mentioned improvement can also be observed in Fig. 6a. Similarly, when they were further split according to the switch and non switch condition (Fig. 6b), this was true as well. The only exception was one subject in the switch condition.

**Figure 5:**
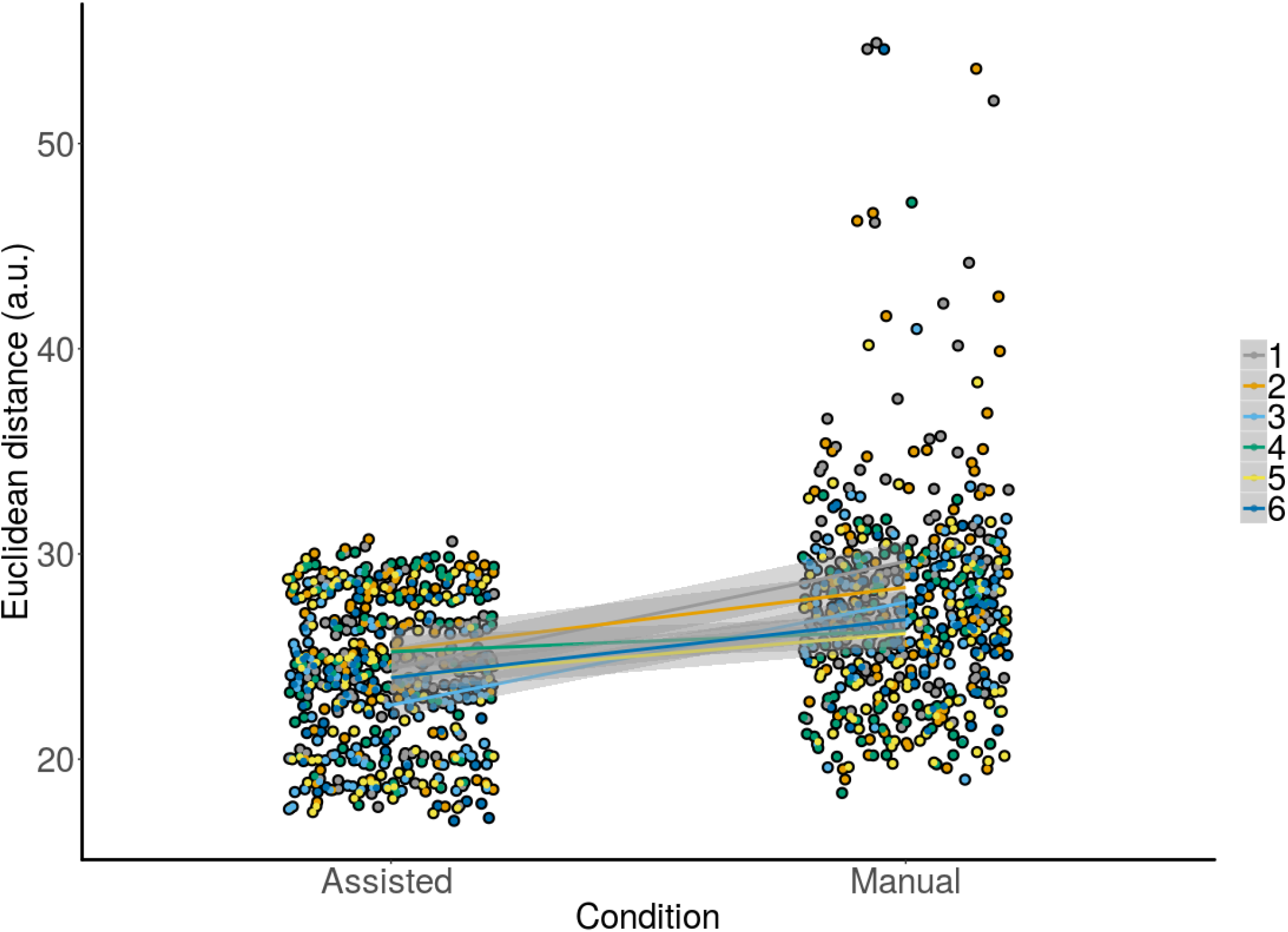
Raw data with corresponding overlaid regression lines and shaded standard errors for the assisted vs. manual condition across all subjects. The y-axis shows accuracy for all trials for each subject individually. A positive slope from the assisted to the manual condition means that our controller improved accuracy across trials. This can be observed for all subjects.

**Figure 6:**
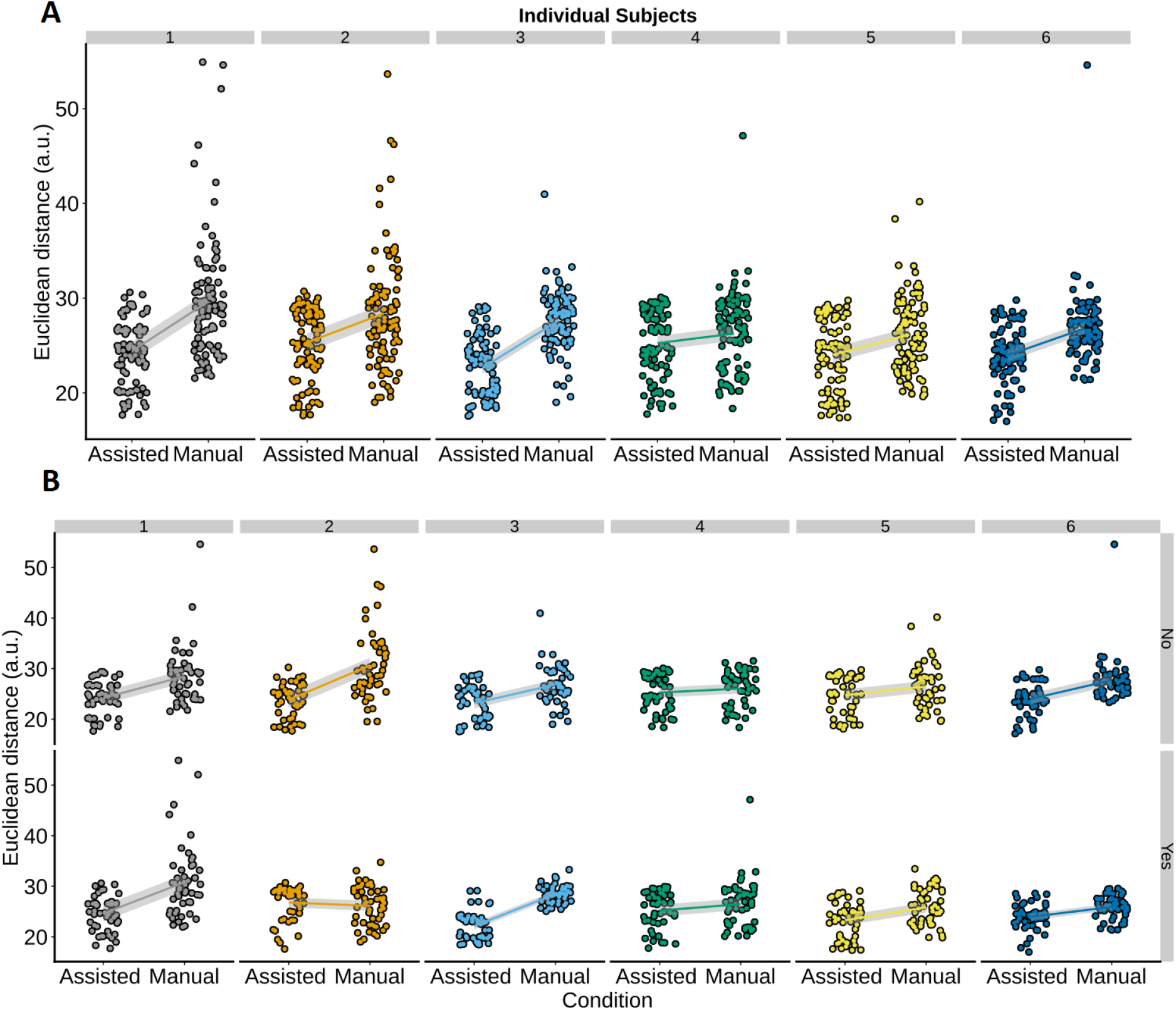
Raw data separated by subject with overlaid regression lines and shaded standard errors for the assisted vs. manual condition. The y-axis shows accuracy for all trials for each subject individually. A positive slope from the assisted to manual condition means that the controller improved subjects’ accuracy. A positive slope can be observed for all subjects in panel **a)** data was collapsed across the switch and the no switch condition. Similarly, when the data was split according to the switch an no switch condition, a positive slope can be observed for all subjects and conditions in **b)**, except for the bottom row of second panel.

In sum, all the described tests that assessed the difference between shared/assisted and manual control provided congruent evidence for our hypothesis. That is, that shared control, where the TV-LQR provided assistance, will enable subjects to be more accurate on designed grasping task compared to manual control where they did not receive any assistance.

As an auxiliary measure, Fig. 8 shows all trials for one subject. Only the *x* and *y* dimensions were plotted for clarity purposes, with Supplementary Materials Fig. 4 showing how the trajectories used throughout the testing looked like. The grey (manual) and orange (assisted) lines show individual trajectories of system poses over a series of trials where the blue line (trajectory I) represented the target trajectory. The target trajectory was picked arbitrarily. The black lines represented the non-target trajectories. Additionally, trajectory D was plotted in red because it was a trajectory close to the target one onto which the TV-LQR controller converged in a small number of trials. For a bundle of runs in the assisted condition, it is noticeable that the algorithm first filtered an incorrect trajectory until approximately 3/4 of the trial, after which it started converging on the correct target trajectory. A similar pattern is observable on a few trials where filtering first favored the incorrect trajectory D but then shifted towards the correct target trajectory I towards the end.

Finally, Fig. 8 also allows for visual inspection of the inaccuracy of the manual compared to the assisted condition, considering that the goal in both conditions was identical. That is, to come as close as possible to the grasp target pose of the blue trajectory.

#### 2.2.3 Target trajectories are more often achieved in the switch condition under assisted control

The last part of the results was related to prediction of intention. Fig. 7 showed a higher convergence towards one of the target trajectories in the case of TV-LQR assistance (upper row, accuracy for non-switch condition: 27.33%, switch condition: 32.33%) as opposed to manual control (bottom row, accuracy for non-switch: 30.33%, switch condition: 30.00%), regardless of whether switching was required or not. Furthermore, in both the manual non-switch (51.67%) and switch (53.67 %) conditions, the proportion of non-target final trajectory states (all trajectories except F, I, J) was higher when compared to both the non-switch (35.33 %) and switch (29.00 %) case of the shared control condition with TV-LQR assistance, indicative of a higher convergence on the final target goal states and better estimation of proximity of the desired target grasp poses, as picked by the subjects. This shows that despite small differences in accuracy between the two conditions, the shared control condition with TV-LQR assistance provided feedback for irrelevant goal states substantially less often in both the non-switch (16.33%) and switch (24.67 %) condition.

**Figure 7:**
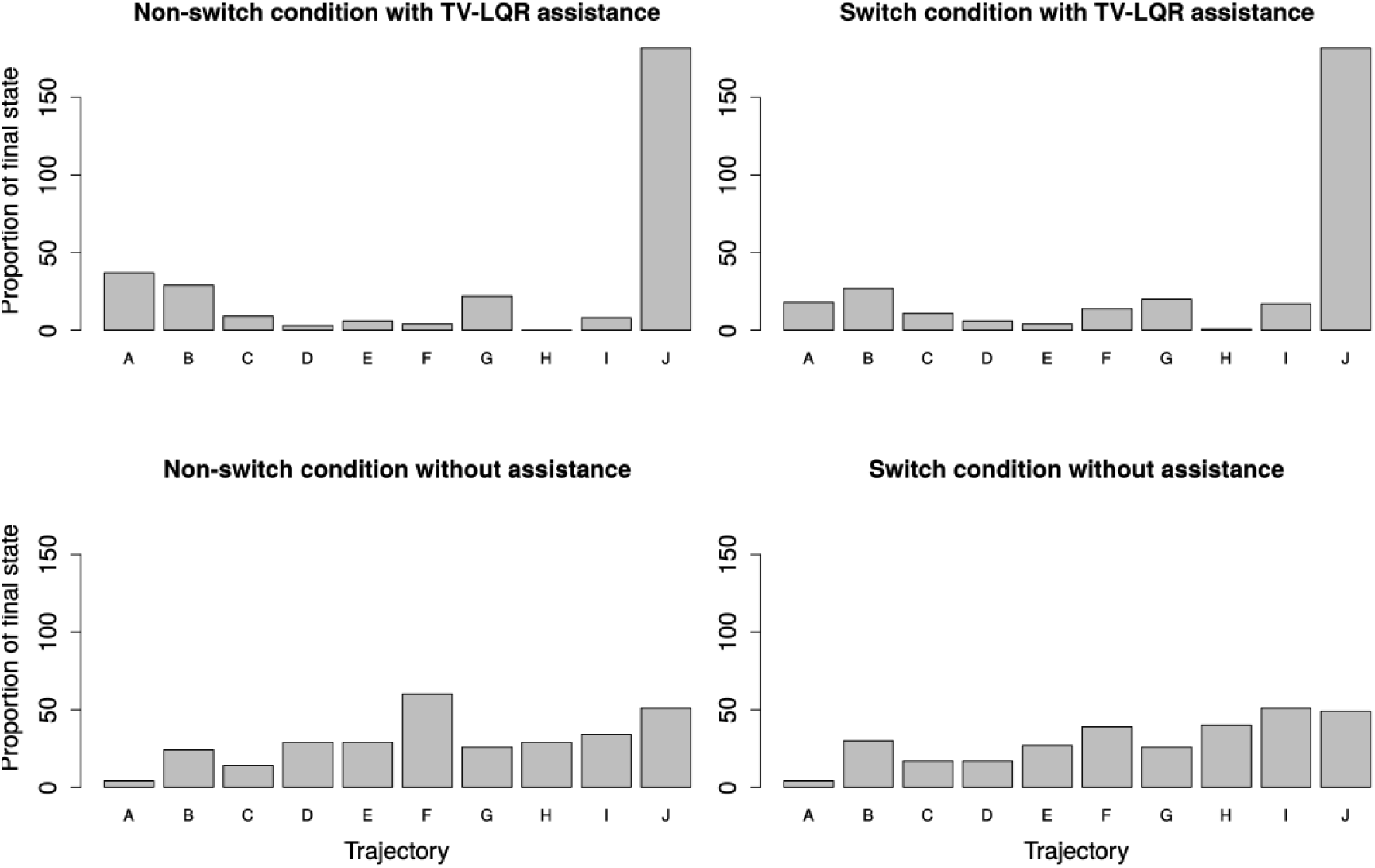
Histogram with frequency of the final system pose being closest to the target grasp pose. In the case of a random visitation frequency expected by chance, each target grasp, denoted by its trajectory name, would be frequented between five and six times (50 trials per condition).

**Figure 8:**
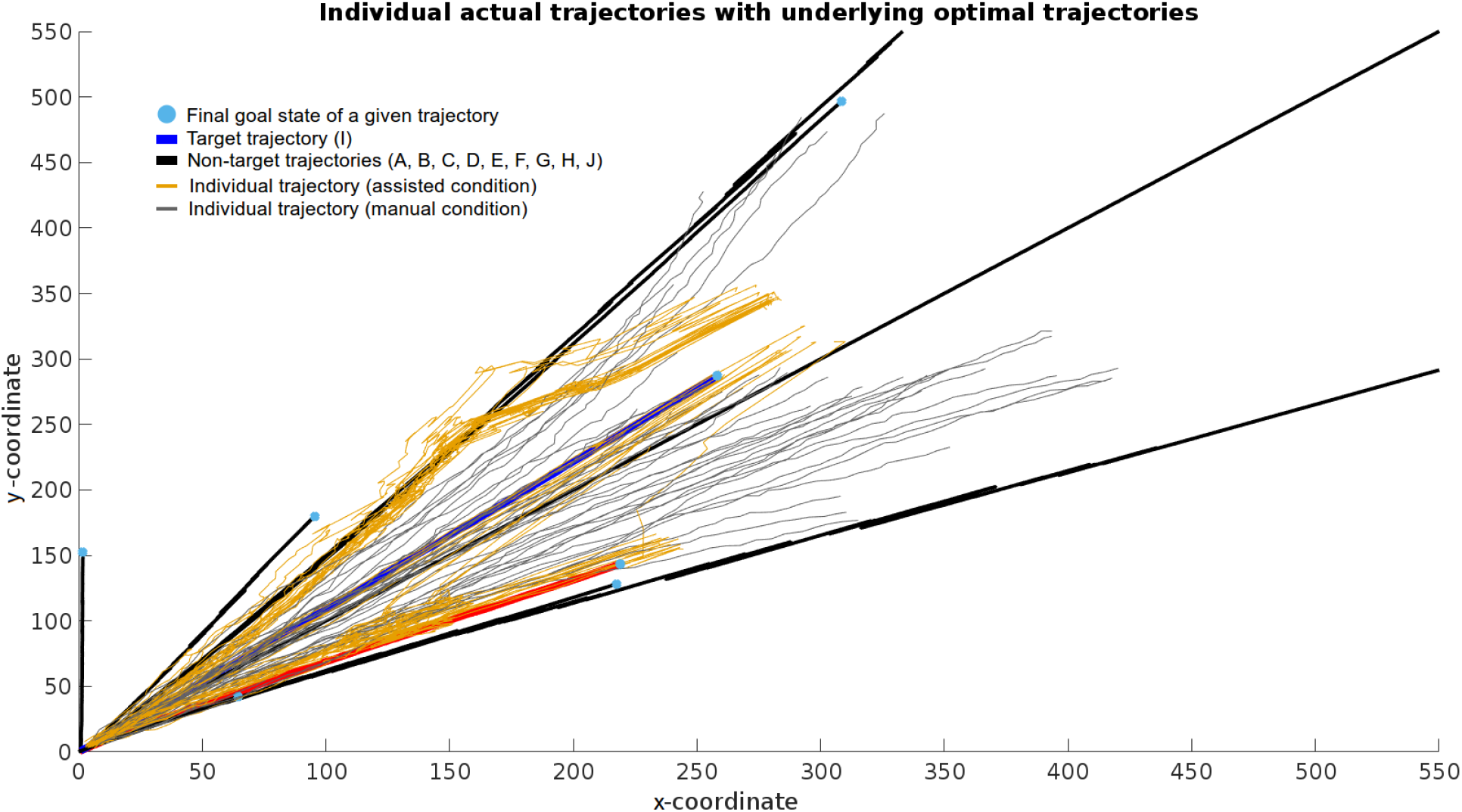
Example data from a series of executions of the manual (gray lines) and assisted (orange lines) condition. Black, red, and blue lines correspond to the optimal trajectories leading to their respective target grasp poses. More specifically, the blue line corresponds to the target trajectory (I), the red line to a neighbouring trajectory (F) to which the assisted condition incorrectly converged in some cases, and the black lines to remaining neighbouring trajectories. This data displays the starting and ending positions together with the error and dynamics of how the controller handled user input from beginning until the end of a given trial.

## 3 Discussion

The presented approach shows how accuracy and thereby usability of semi-autonomous robots for reach-to-grasp tasks, such as human operated manipulators for nuclear waste disposal, can be improved. The key idea of the proposed work is to develop systems that are simultaneously context- and user-aware. Context-awareness was implemented as a black box by manually generating feasible grasp candidates and noisy trajectories which served as input to a TV-LQR controller. In future work, this approach can be extended by building context-awareness automatically. This would allow to employ the proposed system on novel scenes, which has already been shown to be possible in our previous work (*11, 32, 33*). User-awareness was implemented by the filtering that the TV-LQR controller, that was created for all trajectories, performed when the user supplied motion commands.

In brief, the described controller was first tested through characterisation runs. This enabled testing the TV-LQR-based interface while modifying several parameter combinations (*α*, *τ*, *S*) and trajectory lengths described in the methods to assess their impact on accuracy under constant input.

Following the characterisation runs, the TV-LQR was tested on human subjects to determine whether it would improve accuracy compared to manual control. The results showed that the assessed interface as a way of smoothing user input represents a feasible approach for helping to reach goal targets more accurately when compared to manual control. This was shown both in a scenario where subjects were instructed to stay with the same goal from the beginning and where they were required to switch to another goal mid-way.

Moreover, the system was additionally capable of recovering from incorrect predictions, i.e. when the selected target grasp was not the same as the one chosen by the user. This is best observed in Fig. 8, where a bundle of trajectories, which first followed an incorrect optimal reach-to-grasp trajectory, started shifting towards the one the user picked (blue line), and finally converged on that one.

Crucially, because the aim was to build a robust controller system that would mimic real world use cases of remote tele-operations, all the trajectories that were generated had added noise. Similarly, all the tests reported in the results were also performed with non-additive, noisy updating of the system at each waypoint of each trajectory. The main reason for these design decisions was to ensure that the controller would generalize to real world noisy scenarios well, as both would be predicted when observations are made from incomplete information, e.g. in nuclear waste disposal scenarios and subsequent faulty odometry reading that would render such a system less useful.

Furthermore, the experimental setup for the described controller was a 2D simulated scenario for reach-to-grasp tasks (e.g. Fig. 8, Fig. S4). Such a scenario has several advantages for the tests reported in this work. Their simplicity allowed for full control of the environment and a safe test of characterisation runs. Moreover, due to this it was possible to assess the benefits of adding context- and user-awareness with respect to manual control by mimicking a real state-of-the-art setup without potential confounds that could have affected the proposed comparison. Namely, setups in real-world settings require months of training for a human operator to become proficient in a high DOF setting on one or several 2D displays. This is due to the rudimentary and contra-intuitive interfaces employed, i.e. manual control in joint or Cartesian space, and the lack of depth perception provided by a limited number of fixed cameras on the robot site. In contrast, the employed experimental setup circumvented this potential problem to focus and test the proposed framework on naive users after a few training trials. This was due to the fact that the employed experimental setup did not require the users to learn a complex 2D-to-3D mapping between the 2D feedback of a computer display and the 3D environment while providing a full visual description of the scene.

Similarly, the two scenarios to which the subjects were exposed during testing represented pHRI real-world use-cases when a human operator would typically be controlling a manipulator that would need to be moved from a starting pose to a final goal pose along several possible trajectories. The non-switch condition corresponded to the case where the human operator would have a final goal pose in mind (e.g. moving a manipulator to a canister), that would then be followed from the beginning to the end without changes. The alternative scenario, the switch condition, would be one where the operator would change the intended final goal pose at some point throughout the execution, e.g. because of realizing that the intended path is blocked. In this case, the user would need to pick a new target goal pose. The proposed system allows the user to improve the accuracy in reaching the preferred final goal in both scenarios compared to manual control. This work has further shown that that extending the TV-LQR controller with a predictive formulation can be used to infer motor intention in a highdimensional scenario where the TV-LQR assisted condition less often ended up on any of the irrelevant target goal states compared to the manual control and showed worse (non-switch) and better (switch) performance in terms of its accuracy of correctly predicting intended goals. The practical utility of this controller could be further assessed in users who, for example, teleoperate surgical robots or, alternatively, exhibit suboptimal motor control. By doing so, the proposed approach would be further evidenced as a robust and computationally cheap way of meeting the needs of increased pHRI due to automation and advanced controller systems designed for humans with diagnosed motor disorders (*1, 4, 7, 8, 28*). It is worth noting that by assessing it in a complex domain (i.e. with limited or sparse information for the user) this would likely yield better results for the TV-LQR controller. Indeed, it is reasonable to assume that as the complexity of the testing domain increases, the TV-LQR approach would increasingly outperform manual control. At the same time, this represents the main limitation of this work since the trajectories were generated arbitrarily and tested in simulation. However, despite being performed in simulations, due to three sources of noise that were added on long trajectories and the reasoning described above, this proof of principle approach should scale to real-world scenarios. Moreover, further investigation is required to extend this framework to tasks beyond grasping, such as peg-in-the-hole problems when the interactive forces arisen by the physical interaction become critical (*27*).

Overall, the results show that using a TV-LQR with a predictive formulation can be used to improve grasping performance in terms of accuracy together with robustly predicting movement intention and recovering from incorrect predictions in an online fashion. This was achieved in simulation where both the controller and the environment separately included noisy components as a proof of principle test for the proposed controller. Furthermore, the importance of parameter tuning was demonstrated in the first part of the results as an auxiliary component when it comes to optimising such a system. Namely, the employed parameter combination will potentially impact total accuracy of such controllers. This aspect will become increasingly important as personalised controller systems will need to account for interindividual variability of human users in terms of their motor capacity and control characteristics (i.e. the same default parameter combination might not be optimal for every user), a notion that has been long-acknowledged in other fields (e.g. personalised medicine (*24*). In sum, the TV-LQR with a predictive formulation represents a possible component of future workplace, clinical (e.g. motor disorders), or every-day (e.g. exoskeletons for an aging population) pHRI systems where shared control (*16*) of human and machine that jointly exert the environment will be vital.

## 4 Methods

### 4.1 Complete setup

The proposed TV-LQR implementation is part of a robot grasping framework previously described in (*11*). Here we present an important innovation of this framework that enables a human user to control the position of the robot arm using a control input device (e.g. keyboard, joystick, or haptic device) which is filtered in such a way as to always keep the state of the robot arm along the trajectory of the user target goal. In this way, we translate the TV-LQR implementation to a pHRI scenario. When presented with a scene (e.g. a table with objects available for grasping such as a kettle or a bottle), the grasping framework can create optimal trajectories for potentially feasible grasps for the mentioned objects from a given starting pose (i.e. the initial position of the robot arm). It is then able to execute a given trajectory and grasp a specific object. In the current extension, if the scene has a number of graspable objects, the controller would support the user to follow one optimal trajectory based on the prediction of the user intention. Importantly, this extension allows the user to switch the target goal with the controller accordingly exerting less corrective force (Supplementary Materials Fig. 1-2). The continuous corrective feedback from the controller is received until the pose of the system reaches the target goal of the trajectory. At this position, the grasping mechanism implemented within the grasping framework would become active and grasp the target object. The experimental setup aims to demonstrate that our proposed framework can be used to improve reach-to-grasp performance in tele-operated systems by robustly predicting movement intention.

### 4.2 Formal Characterisation of the State-Space Model and TV-LQR

The implementation of the TV-LQR is described in terms of a discrete, state-space model:

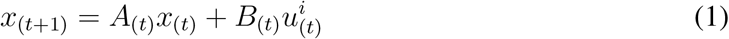

Where *x* represents the system state and *u^i^* represents the system input. All subscripts (i.e. (*t*) and (*t* + 1) represent time notation). In this formulation both *x* and *u^i^* are parameterised as vectors where *x* = [*x, y, z, ϕ, *ψ*, θ*]^T^and 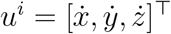. The terms 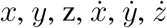 correspond to the coordinate directions and their respective velocities. The angles *ψ*, *ϕ*, *θ* represent rotation for each of the individual axes. The angles were added in the parameterisation for testing how the TV-LQR handles a 6-dimensional system but are not further used. The passive transition dynamics are given by *A*_(*t*)_:

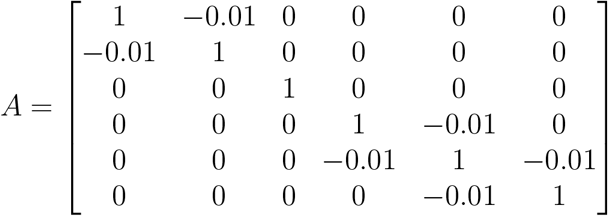

Such a parameterisation leads to a small amount of damping in both the *x* and *y* direction and the **ψ** and *θ* angles. A simplifying assumption that was made was linear dynamics for the *z* direction to make the system continuously move along one of the axes to simulate movement towards the target goals. Finally, *B*_(*t*)_ represents the control matrix, i.e. how strongly will the user input affect the updating of the overall system.

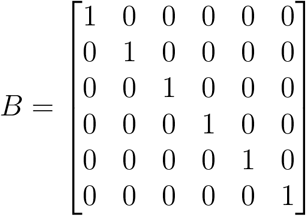

For *B*_(*t*)_ linear dynamics were assumed to avoid modulating the user input and because any non-linear modifications would have been arbitrary in this scenario. For the testing scenario a number of trajectories were generated each with their corresponding number of waypoints. They are defined as the state 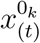 and optimal input 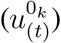 to reach the next state 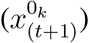 with *k* denoting a potential trajectory. Using the state space formulation above, at each time step the current position (*x*_(*t*)_) and input 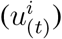 are locally linearized w.r.t. to the trajectory state 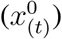 and input 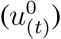 when performing computations:

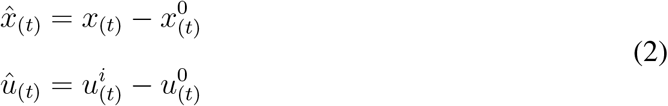

By doing so, the system is reparameterised in terms of the trajectories that are feasible locally at each time step and thus it is not assumed it would stay the same. This step is done for each potential target trajectory.

The obtained estimates from Eq. 2 are used at each waypoint to compute the quadratic cost of the current system state and input:

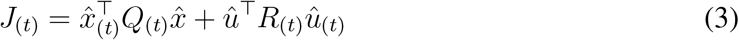

The *Q* and *R* cost matrices are defined as:

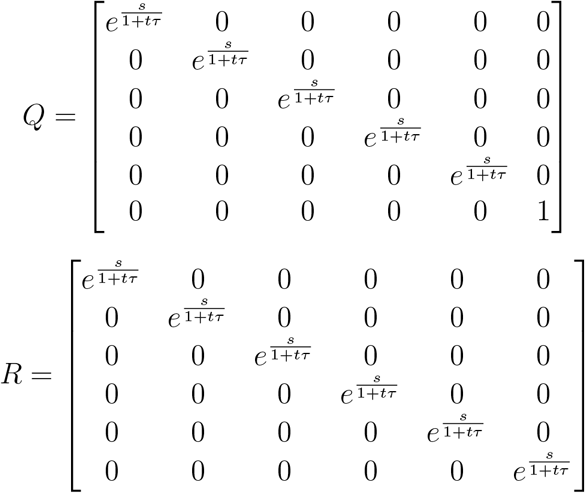

Where the *Q* matrix represents penalisation for being away from the optimal trajectory while the *R* matrix represents how strongly a divergence of the user input against the optimal input is penalised. By minimising the cost function in Eq. 3, the optimal trajectory that is closest to the current system state is found. Importantly, each time before the optimal trajectory can be found, the closest waypoints regardless of the trajectory (i.e. whether the system state is at the beginning or towards the end of its execution cycle) need to be determined first (Eq. 5). Also it is worth emphasising is that the diagonal terms of both matrices are parameterised over time with a hyperbolic discounting function (Supplementary Materials Fig. 1):

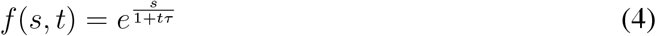

The current waypoint is represented by *t* as an integer (*t* = 1, 2*, …n* = *N*) where N is the final waypoint. The parameter *τ* is a constant reflecting discounting strength of the scaling matrix.

As already mentioned, each time before the optimal trajectory is to be found, the system state needs to have information on the closest waypoints around it to select the trajectory minimising the cost function in Eq. 5. The optimal waypoint is found such that:

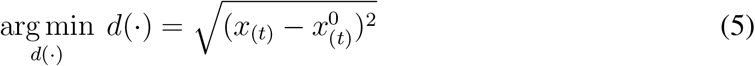

In this case, *d*(·) represents simplified notation for 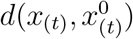 and is the n-dimensional Euclidean distance between the current state *x*_(*t*)_ and states of all trajectories from that waypoint 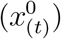 to the final waypoint. Computing the distance from the current sstate to all potential future states of all *k* trajectories was done to remove the assumption of each subsequent waypoint immediately following its predecessor (i.e. this would allow for jumps from waypoint *x* to waypoint *x* + 2 if the user input would have been high enough towards a particular trajectory). Taking together Eq. 3 and Eq. 5, the optimal waypoint of the optimal trajectory with the lowest immediate cost can be selected to make a prediction on where the system state will be at the next time step 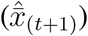:

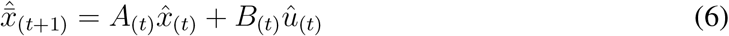

In addition, a conservative estimate heuristic of where the user wants to move next together with a two-waypoint trajectory buffer were added to the formulation. For example, if the user was at waypoint *t* = 15 and the trajectory with the lowest cost was determined to be *K*_(16)_ (i.e. Trajectory K at waypoint *t* = 16), but the last two trajectory waypoints were on trajectory B, the latter was then picked. However, the trajectory with the lowest cost (K) was put in a temporary buffer. Once it happened that in two consecutive observations the lowest cost trajectory was the same and not the one for which the user would later receive feedback, that trajectory was picked (Supplementary Materials Fig. 3).

The utilised heuristic was more important for initial waypoints with a small distance between neighbouring trajectories. Over time it became less important as no crossing appeared in the scenarios presented here. However, to decrease feedback strength when the user wants to switch trajectories, the scheme presented in Supplementary Materials Fig. 2 was employed.

Once the output of the Eq. 6 is obtained, the optimal input 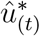 (Eq. 7) can be computed where the term is still formulated w.r.t. to the corresponding 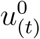.

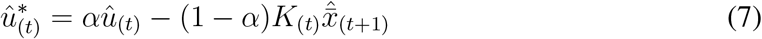

Equation 7 is critical to the updating process, and therefore to the overall system behaviour (i.e. following the nominal trajectory or moving towards a more promising one). It introduces a constant parameter *α*, providing similar functionality to the Kalman gain (*10*). This parameter arbitrates between weighting the pure user input 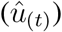 and the state prediction 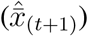 filtered by the feedback matrix (*K*_(*t*)_) (see Fig. 2). This means that high *α* values would lead to strong discounting of the feedback matrix filtering, thereby making the optimal input more dependent on user input. In contrast, low values would lead to strong discounting of the user input and would therefore favour the filtered state information for the optimal input. To compute *K*, the finite-horizon, discrete-case of LQR was used at each waypoint of each trajectory; corresponding to each change of the scaling of the *Q* and *R* matrix

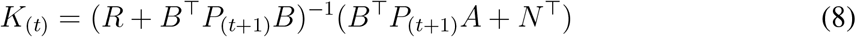

Importantly in the equation above, by computing *P*_(*t*)_ which is obtained by solving the finite-horizon, discrete-case Algebraic Riccati equation (Eq. 9)

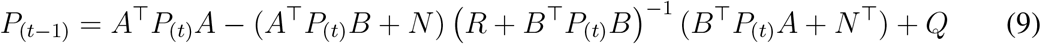

we minimise the cost function:

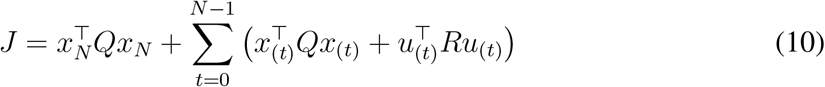

With 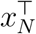 being the final goal state. Finally, the obtained optimal control estimate is dereferenced to obtain the optimal input 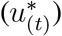 that is then continuously supplied to the system as the control input.

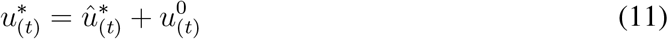

After the optimal input was computed, the prediction estimate of the system pose was obtained using the model from Eq. 1 with a substitution in the input term (Eq. 12) and the corresponding local optimal trajectory state estimate being used in the state term:

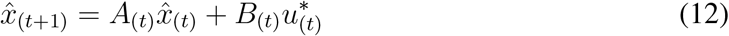

In the last step, the state vector was dereferenced w.r.t. to the local optimal trajectory (Eq. 12) and put back into the world frame.

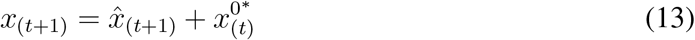

To this state prediction, mean-centred, Gaussian noise with *σ* = 1 was added to both the *x* and *y* coordinate to assess the robustness of our design and mimic measurement error due to either suboptimal odometry or sensor readings (Eq. 13).

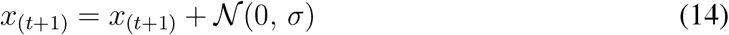

### 4.3 Scenario

The presented system was tested in a scenario where multiple trajectories were generated from an arbitrary starting point representing the starting system pose *x*_0_. The generated trajectories represented potential grasp targets (Supplementary Materials Fig. 4), and were described to the subjects as optimal trajectories from their starting system pose to the potential grasp targets, as outlined in the Procedure. On each trial, each subject was first tasked with providing their desired target grasp (i.e. the name of its corresponding trajectory). This was followed by them continuously providing input to the system by means of a wired computer mouse, with input sampled at 10 Hz. As they provided input, the pose of the system updated and the computations described in the previous section unfolded.

### 4.4 Procedure

The implementation was tested in two parts. The first part is referred to as characterisation runs throughout this manuscript. The second part is referred to as controller tests or testing the controller. In both cases, control was supplied using a wired computer mouse and calibrated on a computer screen using a 1920×1080 resolution, where the coordinate (20, 1020) would represent *u* = [0, 0, 1, 0, 0, 0]^T^. By moving the mouse along the *x* and *y* axis in physical space, one could increase or decrease the velocity of the respective dimension, while the *z* axis would slowly be increased to mimic moving from a starting system pose towards a target grasp pose in one dimension. As described in the Formal Characterisation, rotations were not used in the analysis and due to this the rotation part of *u* was not affected by user input nor modified throughout the tests. Furthermore, in both parts the goal was to go from the aforementioned starting system pose towards a target grasp pose. In the first part, 200 trials were used for each parameter combination to measure their effect on the Euclidean distance between the final system pose and target grasp pose. The amount of trials was estimated to be sufficient for a clear estimate of the Euclidean distance. The model space for the employed was: *τ* (0.1, 0.5, 0.9), *S* (1, 5, 9), and *α* (0.1, 0.5, 0.9). Additionally, trajectory lengths were varied as well with 50, 100, and 200 waypoints through the characterisation runs. A given parameter or employed trajectory length was varied individually while the rest of the test characteristics were kept constant, for each possible parameter combination this was repeated 200 times. Furthermore, Gaussian noise (*σ*) was set to a constant value (*σ* = 1). In general, in characterisation runs constant input was provided by keeping the mouse position (i.e. the velocity) constant beginning with a starting pose until the vicinity of the target grasp pose was reached. At that stage, the Euclidean distance between the final system pose and closest target grasp pose was computed.

While the characterisation results showed that *α* = 0.1 was the optimal parameter for a trajectory length of 200 waypoints, *α* = 0.5 was used to enable equal arbitration between feedback provided by the TV-LQR and user-provided control in the TV-LQR condition (Equation 7).

In the second part, the described setup was tested on six subjects aged between 24 and 27 years with no history of movement disorders and complete upper-limb motor mobility to avoid unwarranted issues in operating the system. Subjects were first familiarised with the system and the input control mechanism described in the previous paragraph. Before starting, they were told they would need to provide input to guide a controller in a simulated 3D scene from a starting system pose to a final system pose that would be as close as possible to one of the target grasp poses, with there being several possible target grasps in the simulated scene, representing possible grasp candidates. Furthermore, they were told there was an optimal trajectory from a given starting pose to a target grasp pose which the TV-LQR would try to keep them as close as possible to in one of the conditions they were about to be tested on. To enable navigation through this simulated 3D scene subjects were presented with information of their current position in terms of *x*, *y*, and *z* coordinates, their velocity, the coordinates of several final target grasp poses, the waypoints of the optimal trajectories closest to their current position, and the coordinates of the trajectory they were actually closest to, in case it was not one of the target ones. In contrast to characterisation runs, the goal of each subject was to pick one of three target grasp poses that were arbitrarily defined at the start of each trial and provide input to move the controller such that they would achieve a final system pose which would be as close as possible to their target grasp pose (i.e. the final pose of a specific trajectory). In controller tests, they were required to switch their targeted grasp half-way through the execution (switch condition) as opposed to the other half, where this was not necessary (non-switch). In both the switch and non-switch condition subjects were required to focus on, arbitrarily chosen, target grasp poses from three trajectories (F, I, J). Therefore, an optimally accurate system would always lead to either of these three trajectories and none of the remaining trajectories (A, B, C, D, E, G, and H). This factor was orthogonal to the assisted and manual condition. That is, to the condition where they were assisted by the TV-LQR when providing input (assisted) and the one where their final system pose dependent entirely on their performance (manual). All subjects were tested on a trajectory length of 200 waypoints and using 100 trials per each of the described conditions, amounting to 200 trials per subject using the optimal parameter estimates from the characterisation runs.

### 4.5 Analysis

The analysis comprised two parts. In the first part, individual conditions of interest from the characterisation tests of the system were compared either using the Wilcoxon rank sum tests or the 2-sided, independent-samples Welch t-tests, depending on whether the parametric assumptions were violated in the comparison. The comparison was done to establish which parameter setting would perform best under constant input. Best was defined as leading to a minimal distance between the final system pose and the nearest grasp target pose. Only accuracy was tested in terms of Euclidean distance, as the speed of execution was not stored nor measured during either the characterisation or main runs with subjects.

In the second part, further Wilcoxon rank sum tests were performed to compare the TV-LQR assisted and manual condition as a whole and segregated on the non-switch and switch condition to assess whether any changes were observed. This provided a direct answer to the hypothesis that the TV-LQR assisted condition would lead to more accurate outcomes. Namely, in the case of higher values in the manual condition, the hypothesis would be confirmed as that would indicate that there was a smaller discrepancy between the final goal state and the system state in cases where corrective feedback was provided by the TV-LQR. As a more stringent criterion, linear linear mixed models were employed to factor in the repeated-measured aspect of the design and note whether the same results would be obtained, i.e. a statistically significant effect of TV-LQR assistance. In addition, for each subject a linear regression on the complete dataset, collapsed across the switch and non-switch condition was performed to assess whether the slope (*β*) coefficient from the assisted to the manual condition would be positive. Namely, this would indicate that for all subjects the same results hold and our results were not driven by outliers (e.g. one subject in the dataset). As an auxiliary component we added Fig. 7 where individual trajectories for both the TV-LQR assisted and manual condition were plotted in a 2D world for visual clarity. These results were not further analysed but were added to provide intuition of the simulation scenario, albeit in reduced dimensionality. Finally, the results were visually inspected to assess whether in the TV-LQR condition, subjects were led to their desired trajectories and how this compared to manual control where no assistance was provided. Here, the final goal states closest to the system pose were plotted as histograms of visitation frequency and checked to assess whether there was a saturation of visitations for the trajectories F, I, and J and whether the pattern of visitations differed between the conditions.

## Acknowledgments

CZ conceived of the presented idea, CZ and SV developed the theory. SV developed the framework, performed and designed the data collection paradigm, the data analyses, and drafted the manuscript. CZ and FD verified the analytical methods. All authors discussed the results and contributed to the final manuscript.

This work was supported by the European Research Council Synergy Project Natural BionicS (contract 810346 to DF).

All data needed to evaluate the conclusions in the paper are present in the paper and/or the Supplementary Materials. Additional data related to this paper may be requested from the authors.

All authors declare that they have no competing interests.

