## Supplements for "Human-Robot Interaction with Robust Prediction of Movement Intention Surpasses Manual Control"

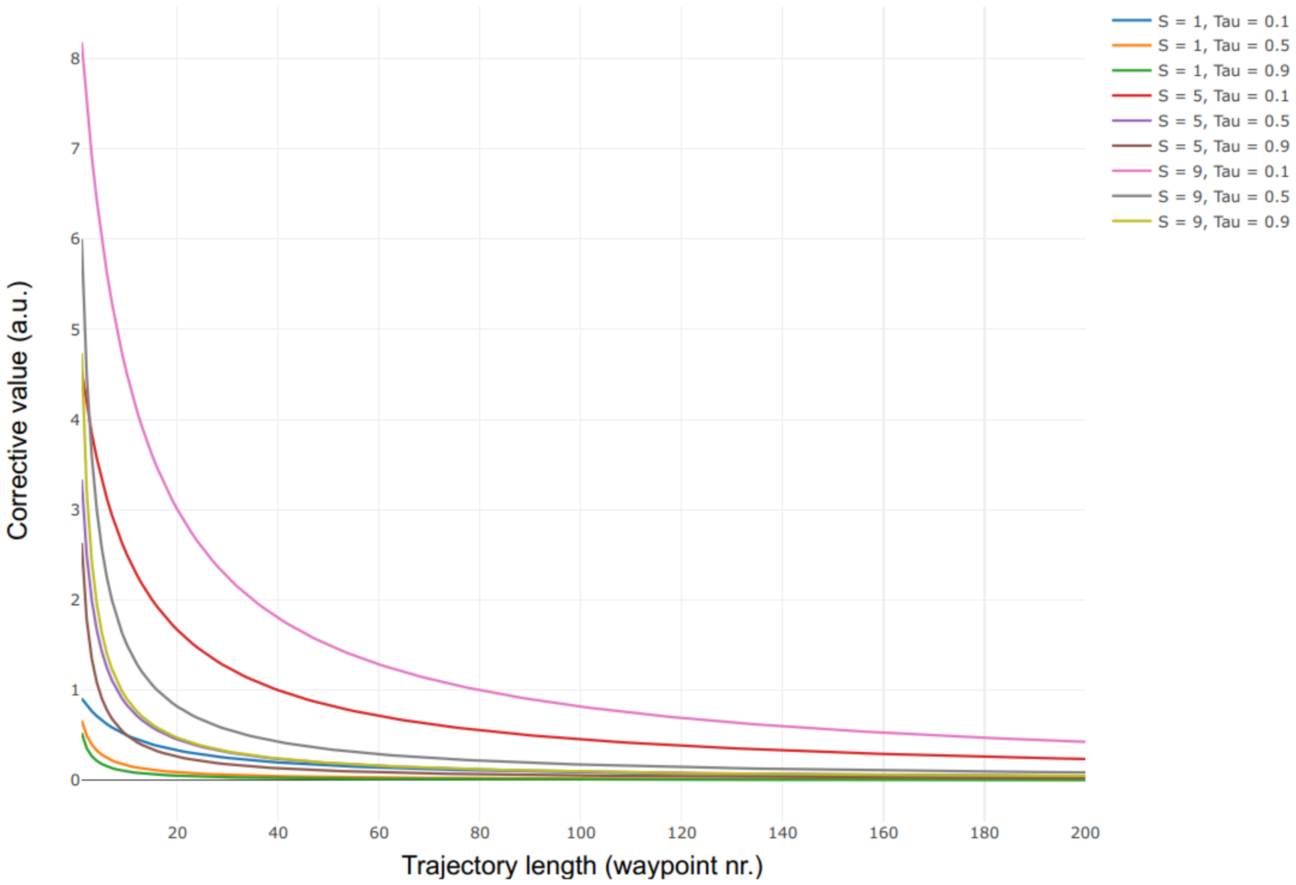


Figure 1: The hyperbolic discounting function used in the Q and R cost matrices on the diagonal elements. $S$ denotes the term in the nominator and is important for the amplitude while $\tau$ denotes the parameter in the denominator and is important for the slope. All parameter combinations are plotted such that the pure effect of the hyperbolic discounter on the Q and R matrices can be clearly observed.

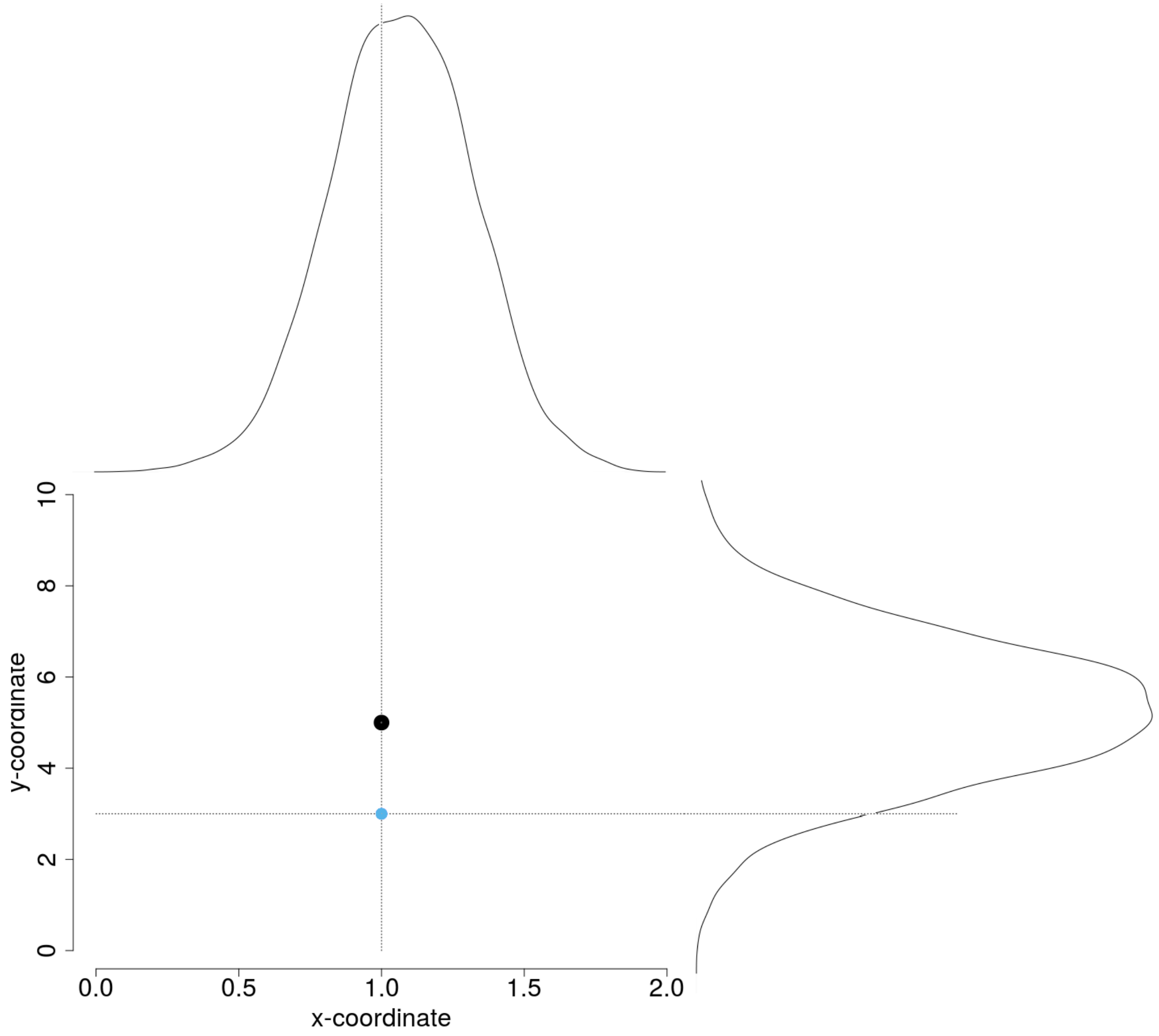


Figure 2: K matrix attenuation. The black dot represents $u^0_{(t)}$ in form of x and y coordinates. The blue dot represents $u_i$. Depending on the proximity of the latter, the originally computed feedback matrix $(K)$ is discounted according to a Gaussian distribution, such that the further away the user is from it, the less force the controller mechanism will exert.

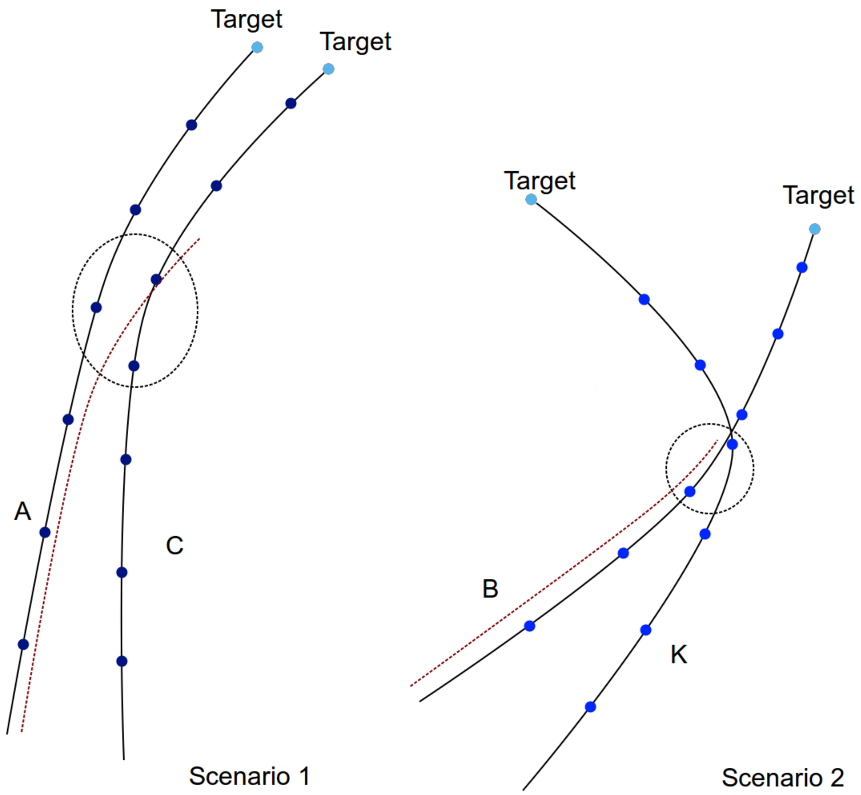


Figure 3: Two scenarios showing the reasoning behind the conservative heuristic estimate. Black lines denote optimal trajectories for the target grasp poses (light blue dots) with discretized waypoints (dark blue dots). The red dotted line corresponds to the actual system pose of a hypothetical user. In the left scenario, the user wanted to switch from trajectory A to trajectory C to move towards another target grasp (target grasp C), as indicated by the red dotted line. Our heuristic allows for sending feedback control for one ”incorrect” waypoint (i.e. feedback control for one waypoint of trajectory A). If it records two consecutive waypoints of the same, new, trajectory (trajectory C), it starts sending feedback control for the new trajectory (trajectory C). A lack of such a mechanism would cause issues in the right scenario. There, due to a crossing between optimal trajectories, the user would obtain feedback control for a trajectory that was not being followed (trajectory K), if our algorithm would always greedily select the optimal solution with respect to distance between the current system pose and the current closest waypoint of a given trajectory.

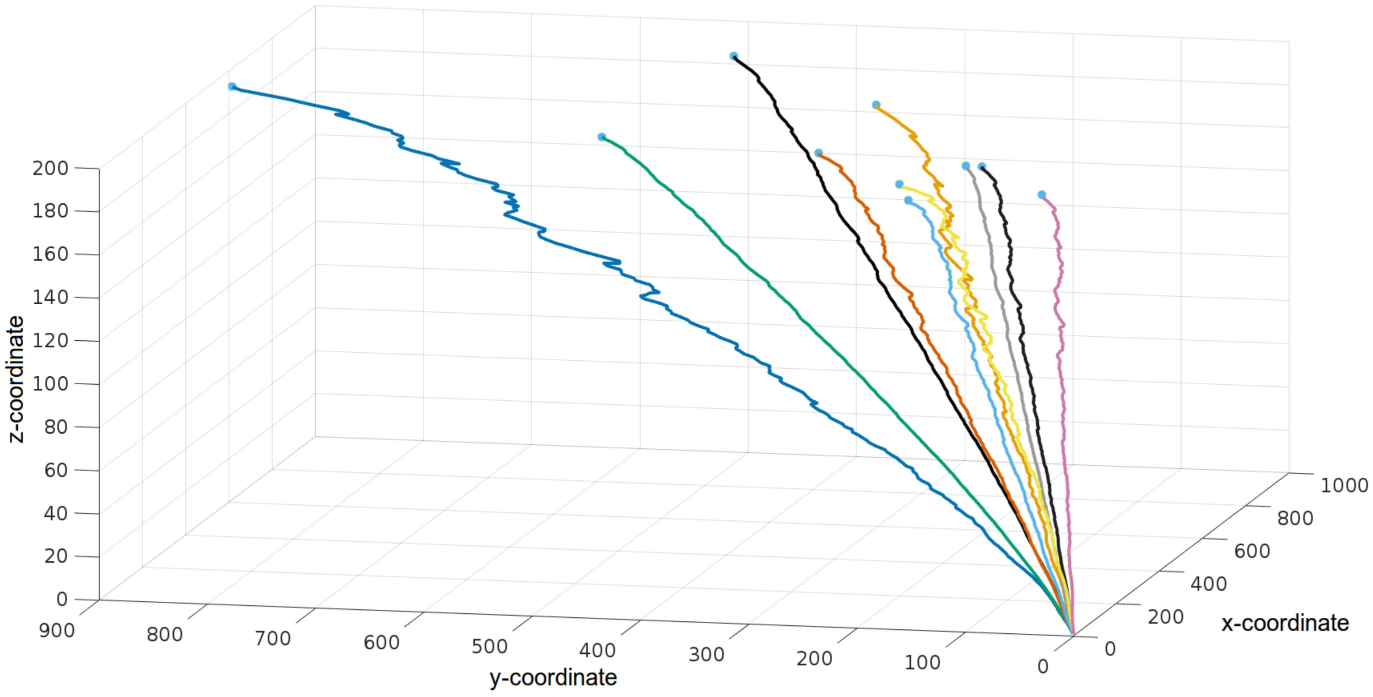


Figure 4: Coordinates of all trajectories used in simulation presented in 3D space. Blue dots on the ends represent goal states for different objects, i.e. the final optimal states for the corresponding trajectories. The individual trajectories show how noisy the testing domain in which the users had to supply input were.
